# The Electromagnetic-Core-First Paradigm: A Mechatronic Co-Design Framework for High-Density Implantable Systems

**DOI:** 10.64898/2026.02.03.703657

**Authors:** Hongyu Li, Yiwen Wang

**Author notes:** **Corresponding author:** Yiwen Wang, **Address:** School of Mechanical and Power Engineering, Harbin University of Science and Technology, Harbin 150080, China., **E-mail(s):**.

## Abstract

The miniaturization of implantable mechatronic systems is fundamentally limited by a pervasive design conflict: within a rigid spatial envelope, the requirements for high electromagnetic power density, physiological fluid pathways, and perfect biocompatibility compete irreconcilably. To resolve this, we propose a paradigm shift—the Electromagnetic Core-First (ECF) co-design framework. Unlike traditional sequential approaches that force performance compromise, ECF establishes a maximized electromagnetic core (via NSGA-III optimization under manufacturing constraints) as the immutable foundation. Fluid and thermal structures are then co-optimized in parallel within this fixed boundary, achieving global system synergy.

Demonstrating its efficacy, we realized a fully magnetically levitated Fontan blood pump (Ø38.2 mm). The ECF-designed prototype, featuring a novel nested magnet topology, achieves: (1) 57.4% higher air-gap flux density vs. baseline designs and an 18.4 N static suspension force at only 4 W; (2) physiological flow matching (8.8 mmHg at 3.2 L/min); and (3) exceptional biocompatibility, with a hemolysis index of 0.0061 g/100 L and surface temperature rise <1.9°C.

This work transcends the presentation of a single device; it establishes a systematic, physics-driven design paradigm that transforms extreme miniaturization from a constraint into a catalyst for performance, applicable to next-generation implantable mechatronic systems.

## 1 Introduction

The relentless drive toward miniaturization in implantable mechatronic systems— encompassing ventricular assist devices, neural interfaces, and targeted drug delivery pumps—has exposed a fundamental and pervasive design bottleneck^[1]^. Within an uncompromising spatial envelope, the synergistic yet competing demands for high electromagnetic power density, anatomically complex fluid or biomechanical interfaces, and perfect biocompatibility create a trilemma that traditional design philosophies struggle to resolve. Conventional approaches, often sequentially prioritizing fluid dynamics or anatomical fit, inevitably force the electromagnetic drive system into a compromised, sub-optimal configuration within the remaining space. This “ design-by-compromise “ paradigm results in a low system-level performance ceiling, limiting operational efficacy, efficiency, and long-term safety in critical applications^[2]^.

This core challenge is acutely manifested in the development of mechanical circulatory support for patients with failing Fontan physiology^[3]^. These patients require a fully implantable, miniature blood pump to restore pulmonary perfusion, posing an extreme instance of the miniaturization trilemma^[4]^: the device must generate sufficient hydraulic power (2–4 L/min at 5–15 mmHg) within a diameter not exceeding 40 mm to fit pediatric anatomy, while simultaneously achieving near-perfect hemocompatibility and thermal safety^[5]^.

Current research predominantly focuses on incremental improvements to axial flow architectures, yet challenges persist. For instance, the Penn State rotary pump^[6]^, while demonstrating promising hemodynamic support, relies on micro-mechanical bearings that raise concerns regarding long-term wear. Similarly, reduced-pressure designs like those by Cleveland et al.^[7]^ and viscous impeller pumps by Yang et al.^[8]^ have achieved compact form factors, but their long-term hemocompatibility profiles remain insufficiently characterized. Consequently, these conventional approaches remain hindered by mechanical bearings or percutaneous drive dependencies, perpetuating risks of thrombosis, hemolysis, and infection^[9]^. The absence of a paradigm-shifting design framework is the critical barrier to a durable, implantable solution.

To break this bottleneck, we propose a fundamental inversion of the design hierarchy: the Electromagnetic Core-First (ECF) Co-Design Framework. This paradigm abandons the sequential compromise model by establishing a maximized, non-negotiable electromagnetic core—comprising the suspension and drive magnets —as the primary optimization objective and fixed physical anchor. Its development is governed by multi-objective optimization (e.g., NSGA-III) that explicitly incorporates manufacturing constraints, defining the Pareto-optimal frontier of force and torque density. Only after this high-performance core is frozen do fluid pathways and thermal management structures undergo parallel, constraint-driven co-optimization within its rigid geometric and electromagnetic boundaries. This “ inside-out “ methodology ensures that peak electromagnetic performance is never sacrificed, transforming the design process from a series of trade-offs into a synergistic integration that pushes the system to its global optimum.

In this work, we articulate the ECF framework as a generalized methodology and demonstrate its transformative potential through the development of a fully magnetically levitated centrifugal Fontan blood pump. The contributions are threefold: We formalize the ECF co-design workflow, detailing its theoretical foundation and implementation strategy for reconciling extreme miniaturization with high performance. (2) We report the realization of a prototype (Ø38.2 mm) whose key innovations—a nested permanent magnet topology for enhanced suspension and a dual-inlet/dual-outlet physiological channel—are direct consequences of the ECF logic. (3) We provide comprehensive multiphysics and in vitro validation, showing the device not only meets clinical hemodynamic targets but also sets new benchmarks for biocompatibility (NIH: 0.0061 g/100L; ΔT < 1.9°C), thereby validating the core hypothesis that prioritizing electromagnetic primacy enables, rather than hinders, system-level excellence.

Beyond presenting a high-performance pump, this study aims to establish ECF as a systematic, physics-driven design paradigm. It marks a critical shift from empirical iteration toward a predictable, model-based systems engineering approach, with broad applicability to the next generation of life-critical miniature implantable devices.

## 2 ECF Co-Design Paradigm and Theoretical Framework

### 2.1 Scaling Law Mismatch in Miniature Systems

To mathematically justify the necessity of the ECF paradigm, this study first analyzes the governing scaling laws for both hydraulic load and electromagnetic capacity with respect to the characteristic dimension (rotor outer diameter, D_out_). Based on the clinical requirements for Fontan circulation, the pump must deliver a specific pressure head (Δ P_*req*_) and flow rate (Q_req_) independent of its physical size. According to Euler’s turbomachinery equation, the pressure head is proportional to the square of the tip speed (*V* = *ωD*_*out*_/2). Consequently, to maintain the required physiological pressure (Δ P_*req*_ ≈ *const*) as D_out_ decreases, the angular velocity $\omega$ must scale inversely:

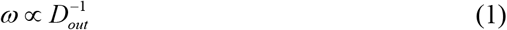

Accordingly, the required hydraulic torque (T_load_), defined as hydraulic power divided by angular velocity (*P*_*hyd*_*/ω*), scales linearly with the diameter:

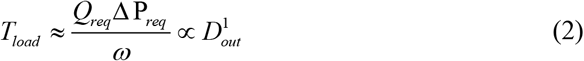

Conversely, for the proposed Axial Flux Permanent Magnet topology, the electromagnetic torque capacity (T_em_) is derived by integrating the Lorentz force shear stress (*σ*_*shear*_) over the annular working surface. By integrating the differential torque from the inner radius (R_in_) to the outer radius (*R*_*out*_ = *D*_*out*_ */*2), and assuming geometric similarity (constant split ratio *λ*) and material limits, the electromagnetic torque capacity is found to exhibit a cubic dependence on diameter:

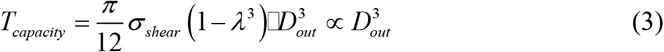

Comparing these two scaling laws reveals a fundamental Scaling Mismatch Inequality:

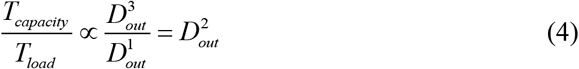

This relationship demonstrates that the design safety factor decays quadratically with miniaturization. As D_out_ shrinks, the electromagnetic capacity (cubic decay) diminishes significantly faster than the fluidic demand (linear decay). This implies the existence of a critical diameter (D_crit_) below which the available electromagnetic volume is physically insufficient. Quantifying this limit by substituting Fontan physiological targets (flow 3.5 L/min, head 10 mmHg) and material limits (N50M magnets, Br=1.4 T) yields a theoretical electromagnetic lower bound of *D*_*en* _ min_ ≈ 34.5 mm. When accounting for rigid manufacturing penalties such as the titanium housing (≥0.5 mm) and fluid gaps (≥0.25 mm), this analysis confirms that for devices below 40 mm, the electromagnetic core is the absolute limiting factor and must be prioritized.

### 2.2 Optimization Problem Formulation

To implement the ECF paradigm, the determination of the electromagnetic core dimensions is formulated as a constrained multi-objective optimization problem. The design variable vector x is selected to define the specific nested magnetic topology. As detailed in Table 1, these variables include the thickness of the magnet rings, the air gap length, and the inner/outer diameters. The search space boundaries were established based on the theoretical scaling limits derived in Section 2.1.

**Table 1:** Design Variables and Optimization Boundaries for Electromagnetic Iron Cores.

| Design Variable | Magnetic Ring Thickness | Air Gap Length | Large Ring Outer Diameter | Large Ring Inner Diameter | Small Ring Outer Diameter | Small Ring Inner Diameter |
| --- | --- | --- | --- | --- | --- | --- |
| Optimization Scope(mm) | 1.1-1.4 | 1.3-1.6 | 30-35 | 35-40 | 17-22 | 22-27 |
| Step (mm) | 0.1 | 0.01 | 0.01 | 0.01 | 0.01 | 0.01 |

The optimization employs the NSGA-III algorithm to simultaneously minimize two conflicting objectives: maximizing the static suspension force (*f*_1_ (*x*) = −*F*_*bias*_) and maximizing the rated drive torque (*f*_2_ (*x*) = −*T*_*rated*_). Mathematically, the problem is defined as:

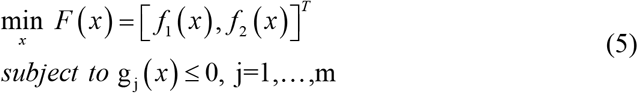

The process is subject to hard constraints to ensure physical feasibility, including geometric validity checks (D_in_<D_out_) and magnetic saturation limits (*B*_*yoke*_ ≤ 1.5*T*).

### 2.3 ECF Execution Strategy and Computational Workflow

Based on the mathematical formulation in Section 2.2, the ECF optimization strategy is executed via a nested co-simulation framework. The core logic of this workflow is to lock the optimal electromagnetic configuration computationally before defining any fluidic geometry. Driven by the NSGA-III algorithm, the process begins by generating an initial population within the bounds of Table 1. For each individual, a logical filter first evaluates the Scaling Limit Constraint (D_out_≥D_crit_). Individuals that violate the fundamental scaling laws are pruned early.

Surviving candidates undergo 3D magnetostatic simulation to compute the objective values, iterating until the population converges onto a stable Pareto frontier. From this converged set, the best compromise solution is identified as the final design choice, representing the optimal marginal utility between suspension capacity and volumetric compactness. Subsequently, the ECF framework enforces a strict ‘Core Freezing’ protocol: the determined vector x_opt_ is established as a set of invariant geometric constraints (г_*core*_). This rigid boundary strictly dictates the spatial envelope available for the downstream stator system and fluid domain integration (elaborated in Section 3).

## 3 Method

### 3.1 Design Objectives and Overall Approach

Guided by the ECF paradigm established in Chapter 2, this section details its implementation in the concrete design of a magnetically levitated Fontan blood pump. The process begins by translating the physiological imperatives of Fontan circulation into system-level engineering specifications, all under the overriding constraint of the theoretical critical diameter (D_crit_) and the scaling mismatch inequality derived in Section 2.1. This translates into two non-negotiable design axioms: (1) Functional Priority: To guarantee robust hemodynamic support and perfect biocompatibility; and (2)Geometric Convergence: To restrict the entire device envelope to within the absolute limit of 40 mm — a boundary not merely imposed by anatomy, but fundamentally dictated by the electromagnetic scaling laws.

Consequently, the verifiable core metrics for the pump are defined as shown in Table 2. The subsequent design workflow is a direct execution of the ECF computational strategy outlined in Section 2.3: first, the electromagnetic core is optimized and frozen; then, the fluid and thermal systems are co-designed within its fixed geometric and physical boundaries.

**Table 2:**
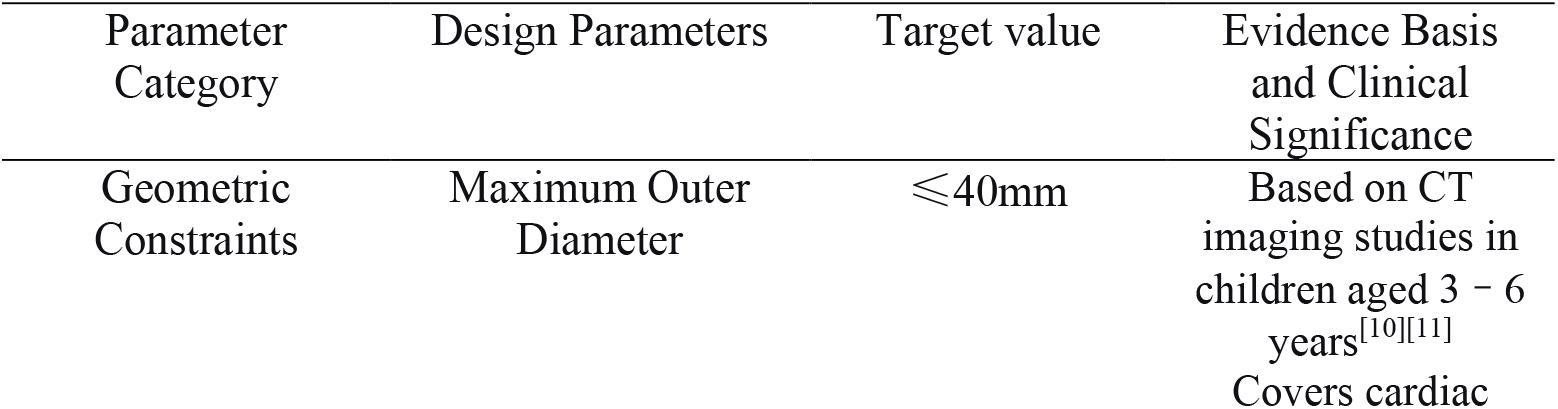
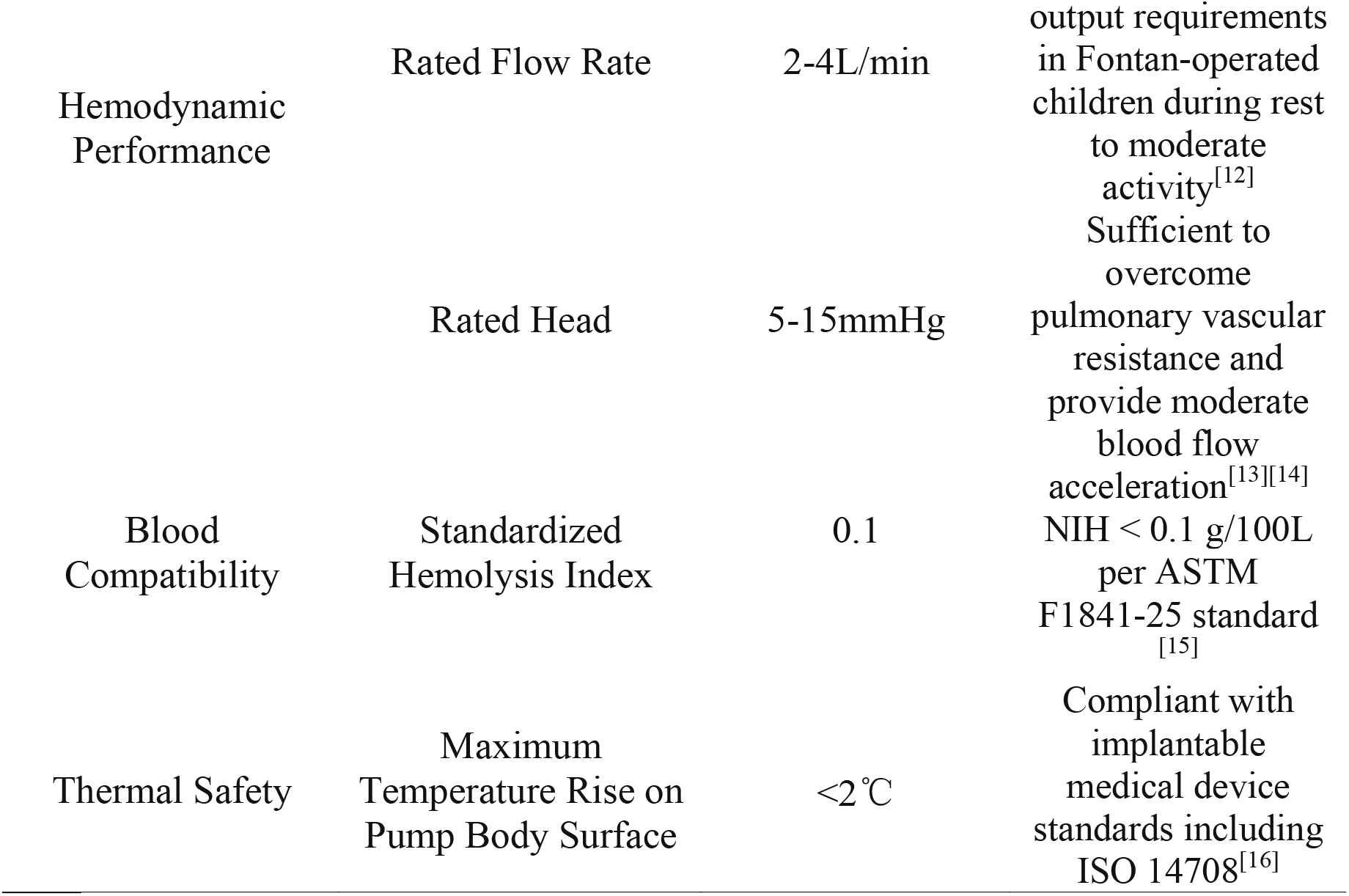
Fontan Blood Pump Design and Performance Targets.

To satisfy these conflicting constraints, we propose an integrated design strategy centered on an “Electromagnetic Core.” The concept prioritizes maximizing the power density of the electromagnetic suspension and drive system within the micro-space, thereby solidifying a high-performance “core.” Subsequently, a dual-inlet/dual-outlet physiological flow path is sculpted around this rigid core, constrained by its geometric boundaries. This “core-first” methodology ensures optimal performance for both the powertrain and fluid systems within an extremely compact footprint (Figure 1).

**Figure 1:**
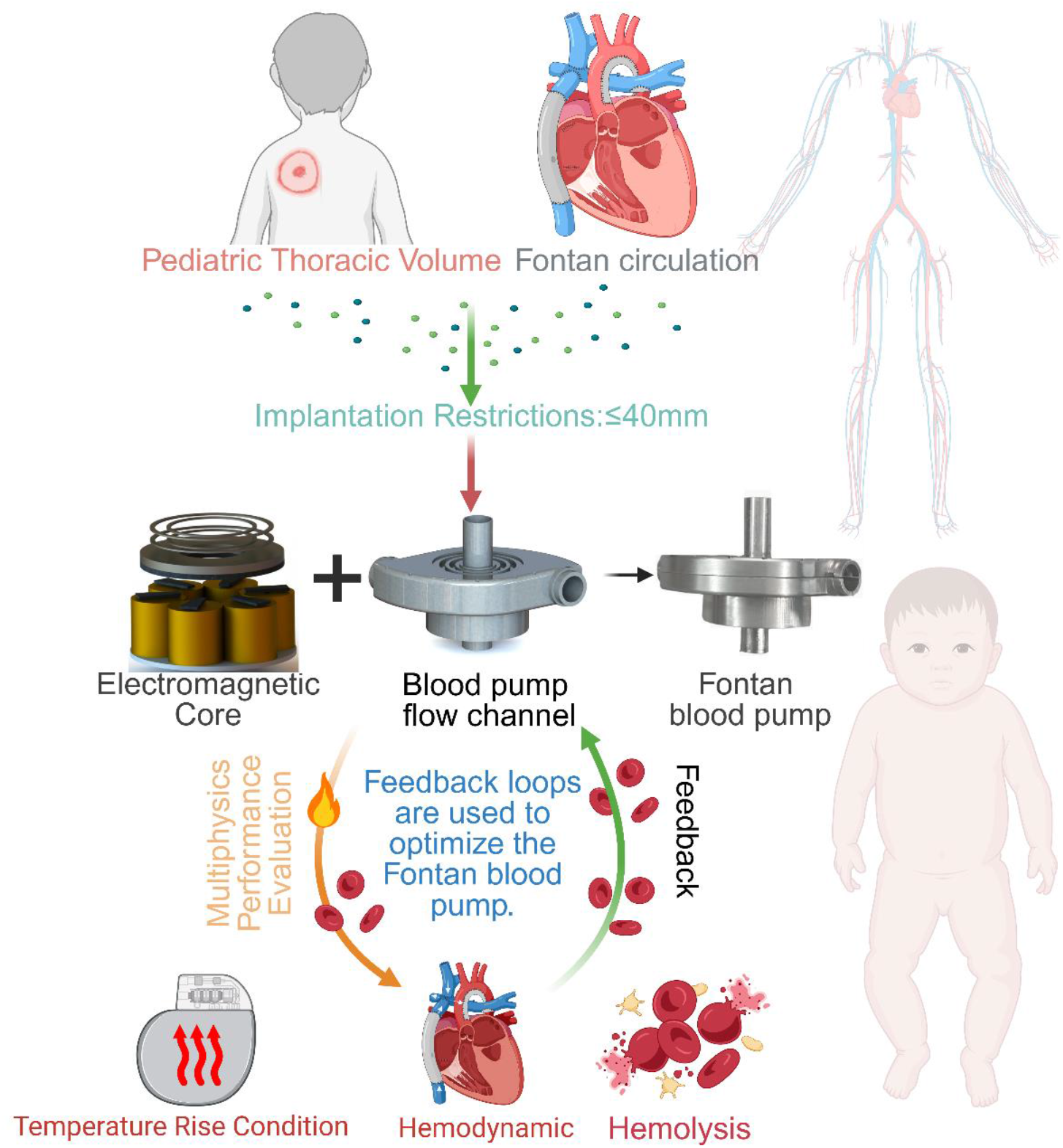
Overall Design and Verification Plan

### 3.2 Electromagnetic Core Topology and Optimization Results

In the ECF methodology, the “electromagnetic core” comprises the rotor-integrated permanent magnet assembly, serving as the immutable physical foundation for subsequent stator design and flow channel integration. Specifically, it includes nested permanent magnet rings for static suspension and a Halbach array for torque generation.

Within the absolute spatial limit of ≤40 mm, a fundamental trade-off exists between suspension load capacity and drive efficiency. To resolve this, we formulated the core design as a multi-objective optimization problem, aiming to simultaneously maximize static suspension force and rated drive torque. The former dictates the suspension margin and static power consumption, while the latter determines pumping capability. Magnetic saturation and volumetric limits were enforced as rigid constraints to ensure physical feasibility.

To achieve contactless operation and miniaturization, a hybrid magnetic levitation scheme was adopted. Permanent magnets provide the primary static bias force to minimize power consumption, while electromagnetic coils exert active control to counteract fluid disturbances, achieving stable five-degree-of-freedom rotor suspension.

A critical innovation lies in the nested permanent magnet ring topology (Figure 2a), designed to maximize force density. Unlike traditional solid disc or single-ring structures that compromise flux efficiency, our nested design synergistically guides magnetic flux through interacting inner and outer rings, concentrating energy within the working air gap (Figure 2b). Finite element analysis (Figure 2c) reveals that under identical constraints (OD 30 mm, Power < 5 W), the nested pair achieves an air gap flux density of 1.1 T—a substantial increase of 57.4% and 50.9% over single-ring (0.85 T) and solid-disc (0.86 T) counterparts, respectively, while reducing flux leakage by 42%.

**Figure 2:**
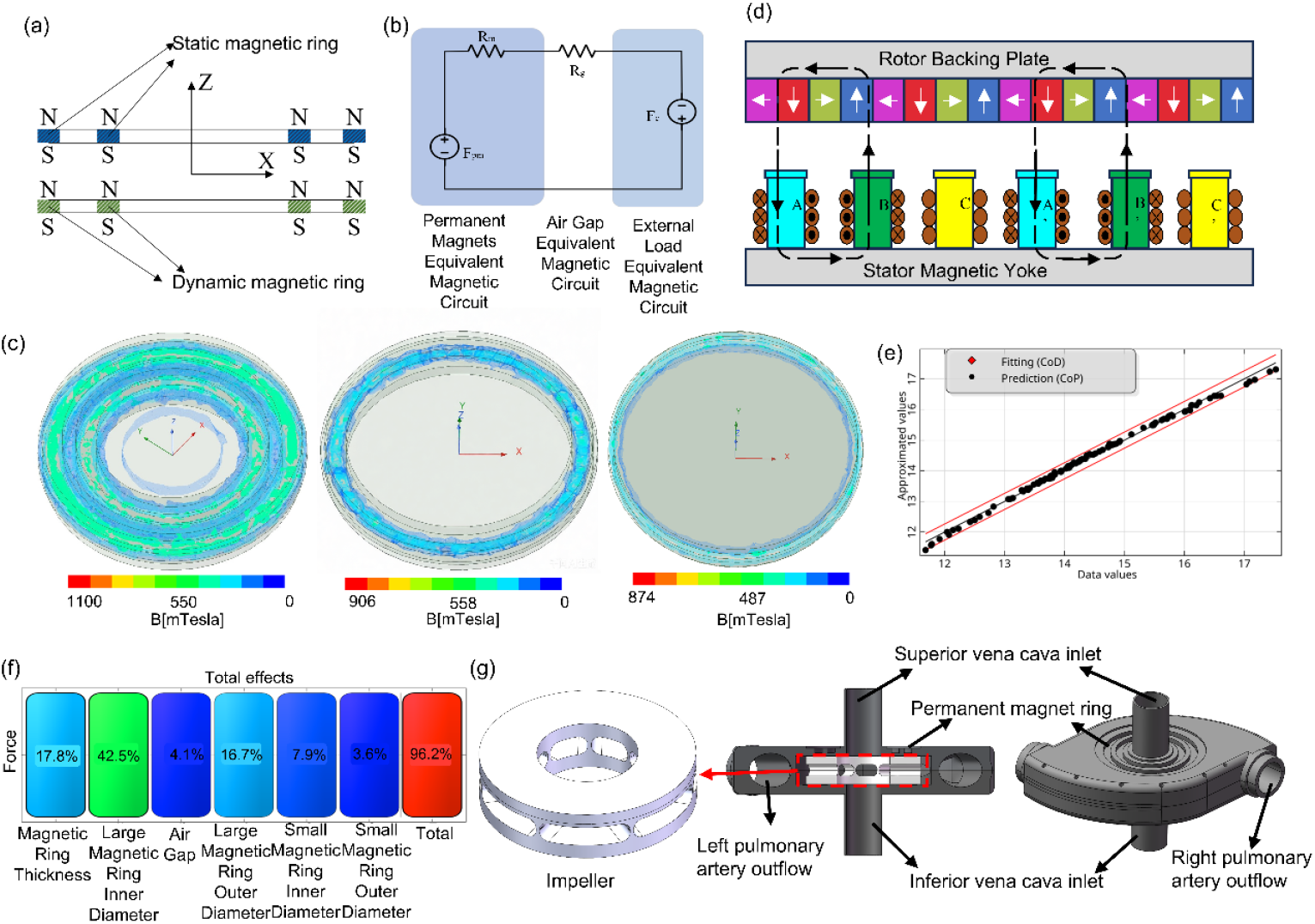
Design Optimization Results for the Blood Pump’s Electromagnetic

Torque generation utilizes a rotor-integrated Halbach array (Figure 2d). Its specific magnetization pattern concentrates a highly sinusoidal field on the air gap side, enhancing torque efficiency, minimizing ripple and stator iron losses, and simplifying stator magnetic circuit design. Given the low-pressure, high-flow nature of Fontan circulation, the motor’s pole pair number (p=4) and Halbach segmentation were optimized for smooth torque output in the low-speed range.

Guided by Fontan hemodynamics (2–4 L/min, 5–15 mmHg; see Table 2), the rated operating point was set at 3000 rpm. Preliminary analysis confirmed this speed provides comprehensive coverage of the target performance window with sufficient control margin for physiological variability, avoiding excessive load increases associated with higher speeds.

Guided by the ECF execution strategy defined in Section 2.3, the geometric parameters for the proposed nested topology were optimized. The specific search space, including parameter ranges and step sizes, is detailed in Table 1. As shown in Figure 2e, the optimization model demonstrates high predictive accuracy relative to actual values. Sensitivity analysis indicates that optimization parameters influence static levitation force and rated torque by 96.2% (Figure 2f), with the most significant factors being the inner diameter of the large magnetic ring, ring thickness, and outer diameter of the large magnetic ring.

The final optimal parameters were determined as follows: ring thickness 1.4 mm, large ring outer/inner radii 14.92 mm/10.30 mm, small ring outer/inner radii 11.07 mm/8.32 mm, and air gap 1.35 mm. Under these conditions, the static suspension force reached 18.4 N.

### 3.3 Stator System Design and Physiological Flow Channel Integration

Once the geometric and electromagnetic parameters of the rotor’s core are frozen via optimization, the design focus shifts to the precise integration of the stator system and the encapsulation of physiological flow channels, strictly adhering to the boundary constraints of the fixed core. The resulting stator-rotor arrangement is illustrated in Figure 2d.

The primary objective of the stator system is to deliver controllable levitation force and efficient driving torque while maximizing electromagnetic coupling with the rotor. An 8-pole/6-slot topology was selected to align with the rotor’s 8-pole permanent magnet structure, generating a synchronous rotating magnetic field. To achieve hybrid suspension and minimize power consumption, segmented permanent magnets were integrated at the tip of each stator tooth. These magnets, in conjunction with the rotor’s nested magnetic rings, form a static bias magnetic circuit that pre-stabilizes the rotor in an axially balanced position. Consequently, the active electromagnetic coils are only required to provide dynamic adjustment forces, significantly reducing control power consumption and heat generation. To predict performance, a lumped-parameter magnetic circuit model was established, representing permanent magnets, air gaps, and windings as magnetomotive force sources and reluctance networks. This model yields analytical expressions for axial suspension force and electromagnetic torque as functions of control current and air gap, providing theoretical guidance for winding parameter design.

The blood flow channel is designed to tightly envelop the fixed electromagnetic core, with its morphology dictated by both the core’s geometry and the physiological pathways of the Fontan circulation. Departing from traditional single-inlet/single-outlet configurations, we propose a novel dual-inlet/dual-outlet centrifugal topology (Figure 2g). Although this complex geometry presents challenges for conventional machining, the curvature continuity was optimized during the design phase to ensure compatibility with high-precision 5-axis CNC machining or additive manufacturing. An integrated molding design was employed to eliminate seal failure risks associated with multi-component assembly. The two inlets align with the superior and inferior venae cavae to enable independent, balanced inflow, while the two outlets correspond to the left and right pulmonary arteries. This anatomical adaptation replicates physiological flow pathways, mitigating abrupt flow turns or recirculation. The impeller blades are positioned centrally within the channel height, creating a symmetrical flow structure that ensures balanced suction at both inlets and shifts operating characteristics toward centrifugal behavior. This yields a smooth head-flow curve suitable for the low-pressure, high-flow Fontan environment. The final device achieves a compact outer diameter of 38.2 mm.

The stator and flow channel designs are not isolated but strictly adhere to a “matching design” principle under the ECF framework. The stator is tailored for optimal electromagnetic coupling with the rotor core, while the flow channel is sculpted to tightly envelop the core for physiological compatibility. This approach establishes a cohesive mechanical structure that exhibits high synergy between core power components and fluid performance.

### 3.4 Multiphysics Simulation and Experimental Evaluation Methodology

Following the 3D modeling of the electromagnetic-structural core and integrated flow channels, a parallel evaluation framework was established to systematically quantify the pump’s hydrodynamic performance, thermal safety, and operational stability. The outputs serve as the quantitative basis for system-level decision-making and design convergence within the co-design process described in Section 2.1.

To primarily evaluate the fluid dynamic performance, a full 3D CFD simulation was employed to quantify pumping capacity and internal flow quality. The fluid domain was extracted from the integrated 3D model, encompassing the complete dual-inlet, impeller channel, and dual-outlet configuration. In ANSYS Fluent, blood was modeled as a Newtonian fluid with a density of 1060 kg/m^3^ and a dynamic viscosity of 0.004 Pa.s. The SST k-ω turbulence model was selected to accurately capture near-wall flow and streamline curvature effects. Boundary conditions were defined as follows: mass flow rates of 0.021 kg/s and 0.0315 kg/s were applied to the two inlets, respectively, and pressure outlet conditions were set according to target physiological operating points. The impeller region was solved using a Moving Reference Frame approach. Steady-state and transient simulations were conducted to verify the pressure head-flow characteristic curve against design targets and to identify potential stagnation zones that could pose thrombogenic risks.

In parallel, thermal safety was assessed via electromagnetic-thermal coupled simulation, which is critical for implantable devices to strictly control temperature rise. Transient electromagnetic simulations in ANSYS Maxwell first calculated copper losses, iron losses, and eddy current losses under rated conditions. These losses were then mapped as volumetric heat sources into a 3D thermal model in the ANSYS Steady-State Thermal module. Boundary conditions mimicked the physiological environment: the inner pump surface contacting flowing blood was assigned a convective heat transfer coefficient of 2000 W/(m^2^.K), while the outer housing surface contacting tissue was assigned 500 W/(m^2^.K)^[17]^. The ambient temperature was fixed at 37°C. This simulation aimed to ensure that the peak surface temperature rise—particularly at tissue interfaces—remained strictly below the 2°C safety threshold.

Finally, to comprehensively validate biocompatibility, in vitro hemolysis experiments were conducted using a customized mock circulation loop (MCL). As high shear stress is a primary driver of blood damage, this physical testing complements the simulation results. Fresh porcine blood, anticoagulated with sodium citrate (9:1 ratio), circulated in a loop precisely maintained at 37°C via a water bath. Blood samples were collected at predetermined intervals during continuous operation. Plasma free hemoglobin (PFH) concentration was measured using a spectrophotometer, and the Normalized Index of Hemolysis (NIH) was calculated based on standard protocols. The quantitative NIH values were further corroborated by qualitative morphological observation of red blood cells (RBC) under optical microscopy.

## 4 Experiments and Discussion

### 4.1 Hydrodynamic Performance: Simulation and In Vitro Validation

To verify that the blood pump meets the preset hydrodynamic requirements, a comprehensive evaluation was conducted combining multiphysics simulation with in vitro experimental validation.

Based on the methodology detailed in Section 2.4, a full 3D hydrodynamic simulation was performed on the optimized pump using a mesh of approximately 3.4 million elements (validated for grid independence). Steady-state results (Figure 3a) reveal excellent flow guidance: blood enters via a streamlined dual-inlet, is efficiently accelerated by the impeller, and exits through the dual-outlet without significant flow separation or recirculation in critical regions. The peak velocity occurs at the impeller tip (∼12.5 m/s); however, given the extremely brief fluid residence time in this region, the shear exposure remains within acceptable limits regarding hemolytic potential. Furthermore, the average outflow velocities at the two outlets are 2.1 m/s and 2.3 m/s, respectively, yielding a flow distribution deviation of <10%, which successfully achieves a physiologically balanced dual-path output.

**Figure 3:**
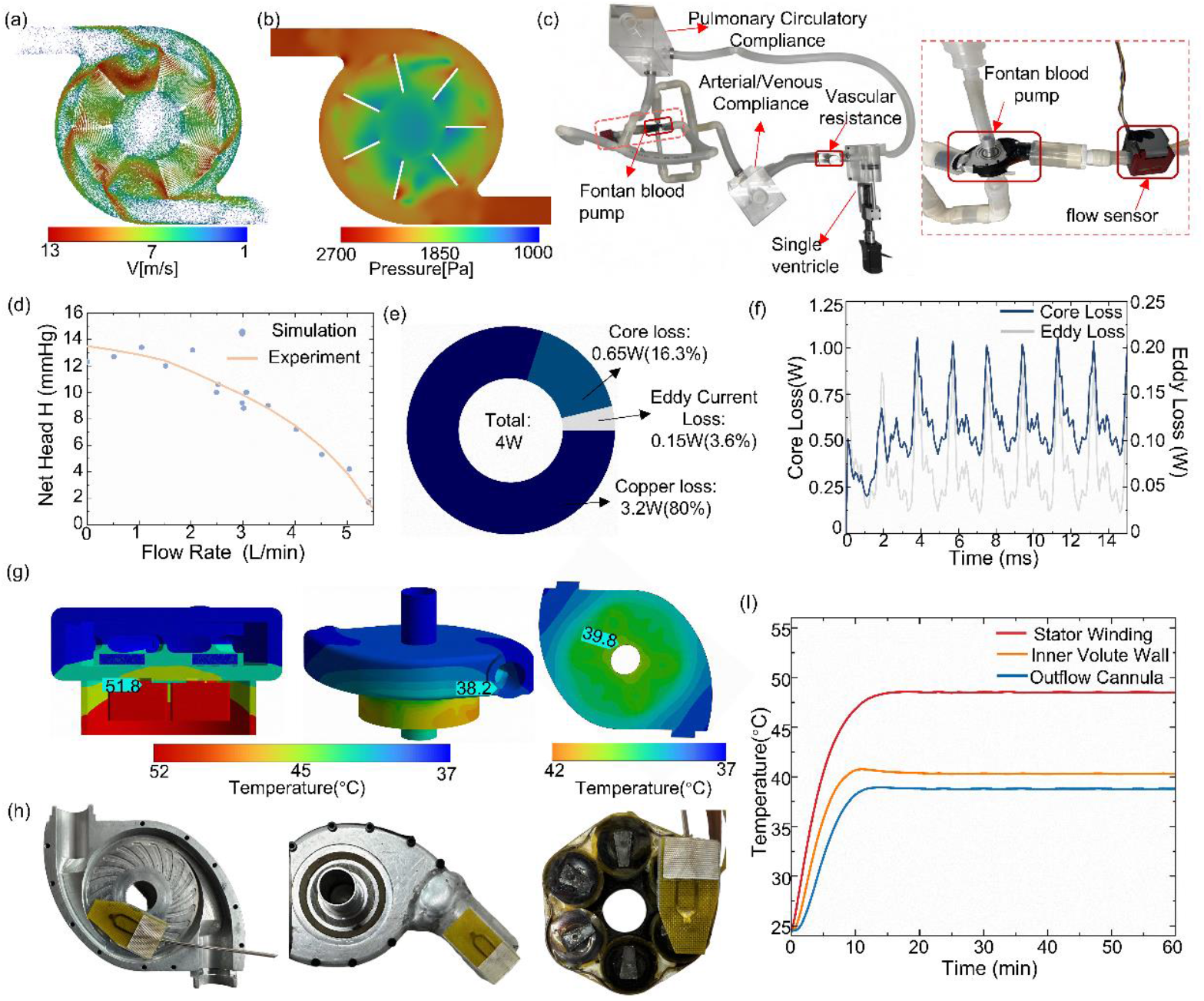
Validation Results of Blood Pump Hydrodynamic Performance and Thermal Safety Performance

Figure 3b illustrates the static pressure distribution at the rated speed (3000 rpm). The pressure exhibits a continuous gradient increase from the impeller center to the periphery, consistent with centrifugal principles. Quantitatively, at the design point of 3.2 L/min, the simulated net head is 8.8 mmHg. This performance precisely targets the mid-flow and upper-head range specified in Table 2, demonstrating that the design not only meets fundamental Fontan support requirements but also possesses sufficient margin to accommodate potential increases in pulmonary vascular resistance.

Prototype Fabrication and Experimental Validation: To validate the ECF design under actual manufacturing constraints, a full-scale prototype was fabricated for hydraulic testing. The housing and impeller were precision-machined from medical-grade titanium alloy (Ti6Al4V) via high-precision 5-axis CNC milling, with geometric tolerances controlled within ±20 μm to realize the complex dual-inlet topology. Blood-contacting surfaces were mechanically polished (Ra < 0.4 μm) and surface-treated to minimize thrombogenic risks.

Hydraulic tests were conducted on a customized MCL using a 40% glycerol aqueous solution to mimic blood viscosity (Figure 3c). Pressure and flow data were collected across the full operating range. Figure 3d compares the experimentally measured H-Q data points with simulation curves. Results demonstrate a high degree of consistency. At the critical validation point (3.2 L/min), multiple experiments yielded an average net head of 9.5 ± 0.6 mmHg. In comparison, the simulation prediction (8.8 mmHg) proved slightly conservative, with a relative deviation of only 7.4%. Notably, the agreement remains excellent from the shut-off point to maximum flow. These results strongly validate the fidelity of the established CFD model and confirm that the manufacturing-aware “ECF” design successfully meets its hydraulic objectives.

### 4.2 Thermal Safety Assessment: Simulation and In Vitro Validation

To ensure long-term implant safety, an electromagnetic-thermal coupled simulation was conducted to evaluate the steady-state temperature rise under rated operating conditions. Electromagnetic losses, derived from the final design in Section 2.2, were mapped as volumetric heat sources. Figure 3e details the loss distribution: with a total power loss of 4 W, copper loss in the windings dominates, accounting for 80% (3.2 W), while core and eddy current losses contribute 16.3% (0.65 W) and 3.6% (0.15 W), respectively. The resulting steady-state temperature contour (Figure 3g) reveals that while the internal stator windings reach a peak of 51.8°C, the surface temperatures relevant to biological safety are effectively controlled. Attributable to the optimized thermal resistance and external convective heat transfer, the fluid domain temperature averages 39.8°C, and the housing surface temperature at the outlet (contacting tissue) is 38.2°C. This corresponds to a surface temperature rise of ΔT = 1.2°C relative to the 37°C body environment, strictly adhering to the <2°C threshold stipulated by implantable medical device standards (e.g., ISO 14708).

To validate these predictions, steady-state thermal experiments were conducted on a customized MCL (Figure 3h). The prototype pump was wrapped in thermal insulation to mimic in vivo tissue conditions and operated continuously under rated physiological load at a constant ambient temperature of 37°C. Once thermal equilibrium was reached (temperature variation < 0.1°C for 30 min), data was recorded via high-precision thermocouples positioned at the stator end, outer housing surface, and inner flow channel wall.

Experimental results (Figure 3i) indicate that under rated conditions, the maximum steady-state temperature on the pump housing (Outflow Cannula) was 38.8°C, while the stator winding end reached 48.5°C. A comparison yields three critical conclusions:

1. Implant Safety (External): The measured external surface temperature (38.8°C) aligns exceptionally well with the simulation (38.2°C), showing a minimal deviation of 0.6°C. The maximum temperature rise (ΔT = 1.8°C) remains strictly below the 2°C clinical safety threshold, confirming no risk of thermal necrosis to surrounding tissues.
2. Hemocompatibility (Internal): The measured inner wall temperature (40.3°C) closely matches the simulation (39.8°C), with an error of only 0.5°C. Crucially, this is well below the 42°C threshold for RBC denaturation, validating safety against thermal hemolysis.
3. Model Robustness: The simulated coil temperature (51.8°C) was slightly higher than the measured value (48.5°C). This indicates that the simulation employed a conservative prediction strategy regarding heat source estimation and boundary conditions. This intentional overestimation enhances the safety margin for actual clinical applications.

In summary, both theoretical and experimental evidence demonstrate that despite generating 4 W of thermal loss, the optimized electromagnetic design and effective thermal management successfully confine potential thermal risks within safe limits, ensuring long-term biocompatibility.

### 4.3 Hemocompatibility Validation: In Vitro Hemolysis Test

Based on the setup described in Section 2.4, a specific loop for hemolysis testing was constructed (Figure 4a). Prior to testing, fresh porcine blood was adjusted to meet ASTM F1841 standards: hematocrit (Hct) to 30 ± 2%, total hemoglobin to < 15 mg/dL, and pH to 7.4^[20]^. These initial values were documented as baselines.

**Figure 4:**
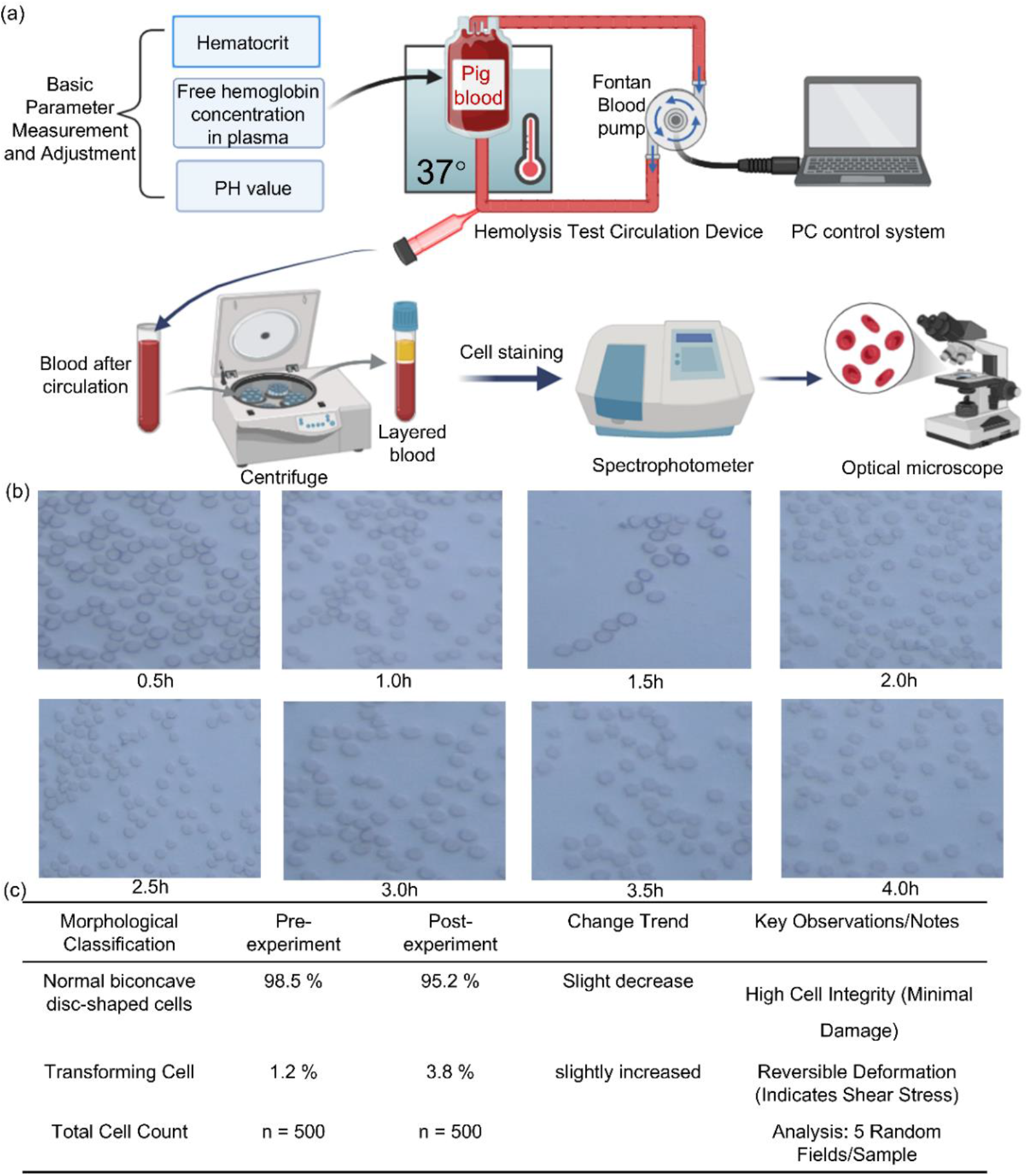
Hemolysis Test Procedure and Results

During the continuous operation of the prototype, blood samples were drawn at 0.5, 1, 1.5, 2, 2.5, 3, 3.5, and 4-hour intervals. Samples were immediately refrigerated to arrest metabolic activity. Plasma was separated from cellular components via centrifugation. Subsequently, PFH concentrations were quantified using a spectrophotometer, while Hct was monitored using an automated blood analyzer. The specific values recorded at each time point are listed in Table 3.

**Table 3:**
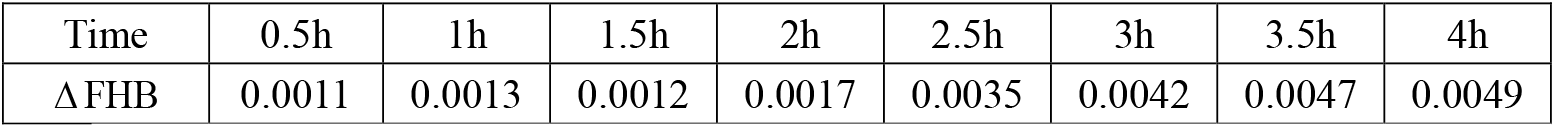
Blood Sample Parameters Collected at Various Time Points.

| Time | 0.5h | 1h | 1.5h | 2h | 2.5h | 3h | 3.5h | 4h |
| --- | --- | --- | --- | --- | --- | --- | --- | --- |
| ΔFHB | 0.0011 | 0.0013 | 0.0012 | 0.0017 | 0.0035 | 0.0042 | 0.0047 | 0.0049 |

Temporal analysis of the results indicates that PFH levels remained low during the first two hours, followed by a gradual increase and subsequent stabilization from the third hour onward. This phenomenon is attributed to two factors: first, the cumulative effect of cellular debris and PFH within the closed loop, which inevitably increases the baseline for subsequent measurements; and second, the system reaching a dynamic equilibrium between mechanical erythrocyte destruction (caused by shear stress or wall adhesion) and the saturation of the fluid medium.

To quantify performance, the NIH was calculated using the standard formula:

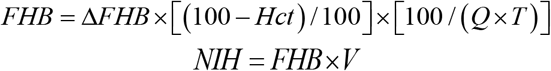

Among these, FHB denotes the free hemoglobin content; ΔFHB represents the incremental value of FHB during the test interval; V indicates the total circulating volume; HCt denotes the Hct of blood (expressed in %); Q is the flow rate of the blood pump (L/min); and T is the time interval (min)^[21][22]^.

The calculated NIH for the ECF-designed pump was 0.0061 g/100 L. This value is an order of magnitude lower than the critical safety threshold of 0.1 g/100 L, confirming exceptional hemocompatibility.

Microscopic analysis further corroborates these findings. Figure 4b illustrates RBC morphology at various operating stages. Post-experiment observations reveal that the vast majority of RBCs retained their healthy biconcave disc shape, with negligible fragmentation. To quantify this, a random sample of 500 cells was manually counted (Figure 4c), revealing a statistically insignificant number of abnormal cells. This confirms that mechanical damage to erythrocytes during high-speed pump operation remains at a minimal level.

In conclusion, the Fontan pump prototype demonstrated superior blood compatibility during the 4-hour extracorporeal circulation test. With an NIH strictly meeting international safety standards and well-preserved cellular morphology, the device validates the effectiveness of the shear-stress-optimized flow channel design.

### 4.4 Discussion

The experimental outcomes collectively affirm the central hypothesis of the ECF paradigm: that in extreme miniaturization, electromagnetic primacy is the critical enabler, not a negotiable constraint. The achieved 18.4 N static suspension force at 4 W exemplifies this, demonstrating that the paradigm successfully circumvents the precipitous decline in force density predicted by scaling laws. This performance benchmark directly validates the theoretical analysis of Section 2.1, confirming that prioritizing the electromagnetic core allows the system to operate far from its fundamental physical limits.

Remarkably, this foundational priority catalyzes advantages across other competing domains. The confinement of the entire device to a 38.2-mm diameter, a direct outcome of the fixed core optimization, imposed a positive geometric discipline on the fluid design. The resulting dual-path architecture, optimized within this immutable boundary, produced a flow field that inherently minimized shear-induced damage. This is conclusively evidenced by the NIH of 0.0061 g/100L, a figure that not only meets but substantially surpasses clinical safety thresholds. Thus, the paradigm resolves a classic design conflict: the pursuit of maximum electromagnetic density inadvertently engineers a fluid environment conducive to exceptional hemocompatibility, by eliminating the geometric compromises that typically create pathogenic flow patterns.

The close agreement between multiphysics predictions and experimental measurements across all modalities—particularly the precise mapping of thermal hotspots and hemolytic regions—underscores the paradigm’s value as a predictive design tool. This fidelity originates from the upfront integration of manufacturing realities into the core optimization loop, ensuring that the transition from digital model to physical prototype is one of high-fidelity realization, not costly rediscovery.

While this in vitro validation establishes a robust proof-of-concept, the natural progression involves confronting the dynamic complexity of the *in vivo* environment. The logical next steps are chronic animal studies to evaluate long-term biocompatibility and system control stability. Nonetheless, the ECF framework’s utility extends decisively beyond this specific application. It provides a generalizable blueprint for the co-design of any high-density implantable mechatronic system—from next-generation pediatric ventricular assist devices to miniature neurostimulators—where the relentless constraints of space, power, and biological integration converge. By redefining the design process from a sequence of trade-offs into a parallel synthesis governed by a physics-driven hierarchy, this work points toward a more systematic and predictable discipline for building the next generation of life-sustaining microdevices.

## 5 Conclusion and Prospect

### 5.1 Conclusion

This study establishes and implements an ECF collaborative design paradigm, providing a systematic solution for developing implantable blood pumps under extreme dimensional constraints. By bridging theoretical rigor with engineering feasibility, this work fundamentally redefines blood pump design: shifting from a “multi-objective compromise” dilemma to a global optimization problem, driven by a fixed electromagnetic core with fluid dynamics and physiological requirements as collaborative objectives.

The core outcomes are summarized as follows:

1. Methodological Innovation: An “inside-out” design workflow was successfully constructed, establishing electromagnetic performance as a fixed boundary. This enables the parallel, synergistic optimization of electromechanical and fluidic characteristics, replacing traditional sequential trade-offs and establishing a generalizable framework for future miniature implantable devices.
2. Engineering Realization: Guided by the ECF paradigm, a fully magnetically levitated micro-blood pump (diameter ≤ 38.2 mm) was realized. This turns miniaturization from a constraint into a performance advantage. By incorporating manufacturing constraints early in the design, the prototype achieved high-fidelity reproduction of theoretical performance, validating the “parallel synergy” design strategy.
3. Performance Validation: In vitro experiments confirm that the device meets clinical hemodynamic requirements while maintaining hemolytic and thermal risks strictly below industry benchmarks. This validates the core hypothesis of the ECF paradigm: prioritizing a rigid electromagnetic core not only avoids compromising fluid performance but achieves system-level gains through global synergy.

Broadly, the value of this work extends beyond the delivery of a high-performance Fontan pump prototype. It demonstrates a systematic, physics-driven design philosophy applicable to next-generation miniaturized implantable medical devices. This marks a critical shift in the field—moving from empirical, local optimization toward a predictable, model-driven systems engineering paradigm enabled by global co-design.

### 5.2 Prospect

Building upon the validated ECF design paradigm, subsequent research will advance along three strategic axes:

1. Clinical Translation and Intelligent Control: Chronic in vivo studies (≥ 6 months) in porcine models will be conducted to comprehensively evaluate long-term hemodynamic efficacy, biocompatibility, and reliability. Concurrently, intelligent closed-loop control algorithms, driven by physiological signal feedback, will be developed to achieve adaptive, personalized circulatory support.
2. Intelligent Design and Manufacturing Platform: To drastically accelerate development, we will integrate machine learning surrogate models with the ECF-NSGA-III framework to construct an automated design optimization platform. This will be combined with high-precision additive manufacturing techniques, such as Selective Laser Melting (SLM), to enable the monolithic fabrication of complex microfluidic structures, pushing the boundaries of miniaturization, performance, and biosafety.
3. Platform Generalizability: Leveraging its core strength in reconciling “high-performance, miniaturization, and biosafety,” the ECF paradigm will be extended to other critical domains. Priority targets include next-generation pediatric ventricular assist devices (requiring diameters ≤ 30 mm and hemolysis indices < 0.01 g/100L) and minimally invasive neurostimulation systems, thereby driving the evolution of implantable devices toward greater intelligence and long-term reliability.

